# The *de novo* genome of the “Spanish” slug *Arion vulgaris* Moquin-Tandon, 1855 (Gastropoda: Panpulmonata): massive expansion of transposable elements in a major pest species

**DOI:** 10.1101/2020.11.30.403303

**Authors:** Zeyuan Chen, Özgül Doğan, Nadège Guiglielmoni, Anne Guichard, Michael Schrödl

## Abstract

**Background:** The “Spanish” slug, *Arion vulgaris* Moquin-Tandon, 1855, is considered to be among the 100 worst pest species in Europe. It is common and invasive to at least northern and eastern parts of Europe, probably benefitting from climate change and the modern human lifestyle. The origin and expansion of this species, the mechanisms behind its outstanding adaptive success and ability to outcompete other land slugs are worth to be explored on a genomic level. However, a high-quality chromosome-level genome is still lacking.

**Findings:** The final assembly of *A. vulgaris* was obtained by combining short reads, linked reads, Nanopore long reads, and Hi-C data. The genome assembly size is 1.54 Gb with a contig N50 length of 8.6 Mb. We found a recent expansion of transposable elements (TEs) which results in repetitive sequences accounting for more than 75% of the *A. vulgaris* genome, which is the highest among all known gastropod species. We identified 32,518 protein coding genes, and 2,763 species specific genes were functionally enriched in response to stimuli, nervous system and reproduction. With 1,237 single-copy orthologs from *A. vulgaris* and other related mollusks with whole-genome data available, we reconstructed the phylogenetic relationships of gastropods and estimated the divergence time of stylommatophoran land snails (*Achatina*) and *Arion* slugs at around 126 million years ago, and confirmed the whole genome duplication event shared by them.

**Conclusions:** To our knowledge, the *A. vulgaris* genome is the first land slug genome assembly published to date. The high-quality genomic data will provide valuable genetic resources for further phylogeographic studies of *A. vulgaris* origin and expansion, invasiveness, as well as molluscan aquatic-land transition and shell formation.

## Background

Land slugs and snails (Mollusca: Gastropoda), which can be found easily in gardens, forests, fields or orchards, in their majority are stylommatophoran pulmonates.

Radiated into about 30,000 species, and highly successfully colonized habitats from polar regions to the tropics, they are among the most successful taxa which achieved aquatic-terrestrial transition [1–5]. Some of them are well-known invasive species/pests across the world. In recent years, the notorious “Spanish” slug, *Arion vulgaris* Moquin-Tandon, 1855, has attracted widespread attention due to its invasiveness and negative impact on economy, ecology, health and social system [6]. As a major defoliator of plants, *A. vulgaris* causes serious damage in orchard cultivation, gardens and agriculture resulting in financial losses [7–9]; transmits plant pathogens, contaminates silage and might cause health problems in animals [10, 11]; it also outcompetes native slug species and reduces the biodiversity [12]. *Arion vulgaris* is considered as one of the 100 worst alien species in Europe by DAISIE (Delivering Alien invasive species inventoried for Europe) and is the only land gastropod among them [13]. The native range of *A. vulgaris* is not known for certain. According to the record of first discovery in many European countries, it was believed that the slug originated on the Iberian Peninsula and expanded its range into central and eastern Europe over the last five decades [6]. However, the very similar external appearance with other closely related native large arionids as well as hybrid species between *A. vulgaris*, *A. ater* and *A. rufus* [14–16], might cause the misidentification of *A. vulgaris*, obscure the specimen records, and make it difficult to trace its origin and monitoring the spread only by morphological identification [17, 18]. Controversially, recent studies based on the genetic diversity patterns of mitochondrial and nuclear loci suggested that *A. vulgaris* is native in central Europe rather than alien/invasive while probably invasive in other parts of Europe [19–21]. In either case, it is undeniable that its outstanding adaptive success, the mass occurrences and consequent pest in the last 40-50 years and the reason behind is worth exploring.

In recent years, genome research has shown its advantages and potential in revealing evolutionary and adaptive mechanisms. However, so far only two land snail genomes were published [22, 23]. Compared to land snails, land slugs have ‘sacrificed’ the protective function of a calcareous shell for less weight and energy costs, less dependence on calcium uptake, better mobility, fast body movements and ability to occupy very small spaces [5]. As a tradeoff, it has to pay a certain price for resisting external stimuli, predators, sun exposure, and drought – or compensate these functions by innovations, e.g., defense by chemical compounds or behavior. However, the genetic mechanisms and evolutionary significance of shell gain and loss has not been explored yet.

Here, we assembled and annotated a chromosome-level genome of *A. vulgaris*, which could provide crucial resources to infer its evolutionary history, elucidate its distribution pattern and explore genetic mechanisms that might be related to its quick adaption and invasiveness. This genetic resource can be fundamental for several lines of applied research, such as for exploring mucus related genes and pathways for medical research [24]. Moreover, the genome is the first land slug genome published to date: by comparison with land snails, it will be a good reference for exploring the evolution of shell losses in Stylommatophora, and potential substitutes of the shell functions, such as chemical weapons or an enhanced immune system in land slugs. Furthermore, the *A. vulgaris* genome will also contribute to molluscan genomics and for studying evolutionary trajectories, e.g., the transition from marine to terrestrial habitats [25].

## Data Description

### Sample collection and sequencing

An adult *A. vulgaris* was collected in the garden of the Zoologische Staatssammlung München, Germany **(Fig 1a)**. Genomic DNA was extracted from the foot muscle tissue with the CTAB method and quality was checked using agarose gel electrophoresis [26]. Four different sequencing technologies were used to obtain the genome sequence. First, one Illumina paired-end sequencing library was generated following the manufacturer’s standard protocol (Illumina) with an insert size of 350 bp. Also, high molecular weight DNA was separated and loaded onto the 10X Genomics Chromium microfluidics controller for barcoding and generated two 10X Genomics linked-read library with an insert size of 350 bp. Those reads not only provided the long-range positional information to assemble contigs into scaffolds [27] but were also used for the genome survey analysis and final base-level genome sequence correction. One Hi-C library digested with MboI and with an insert size of 350 bp was constructed for providing long-range information on the grouping and linear organization of sequences along entire chromosomes to assemble the scaffolds into chromosome-level scaffolds [28]. All these libraries were sequenced on an Illumina HiSeqX Ten platform (Illumina, San Diego, CA, USA) to 150 bp paired-end reads. The raw reads were further filtered with the following criteria: reads with adapters, reads with N bases more than 5%, and reads with more than 65% of low-quality bases (≤ seven) using Fastp [29], which yielded approximately 57 Gb Illumina short reads, 138 Gb barcoded reads and 135 Gb Hi-C reads separately (**Table S1**). Meanwhile, Nanopore libraries were prepared and sequenced in the platform Nanopore PromethION (Oxford Nanopore). 75 Gb long reads with mean and N50 lengths 19,393 bases and 25,802 bases respectively were generated (**Table S1**). Those reads were used altogether for reference genome construction.

**Figure 1.**
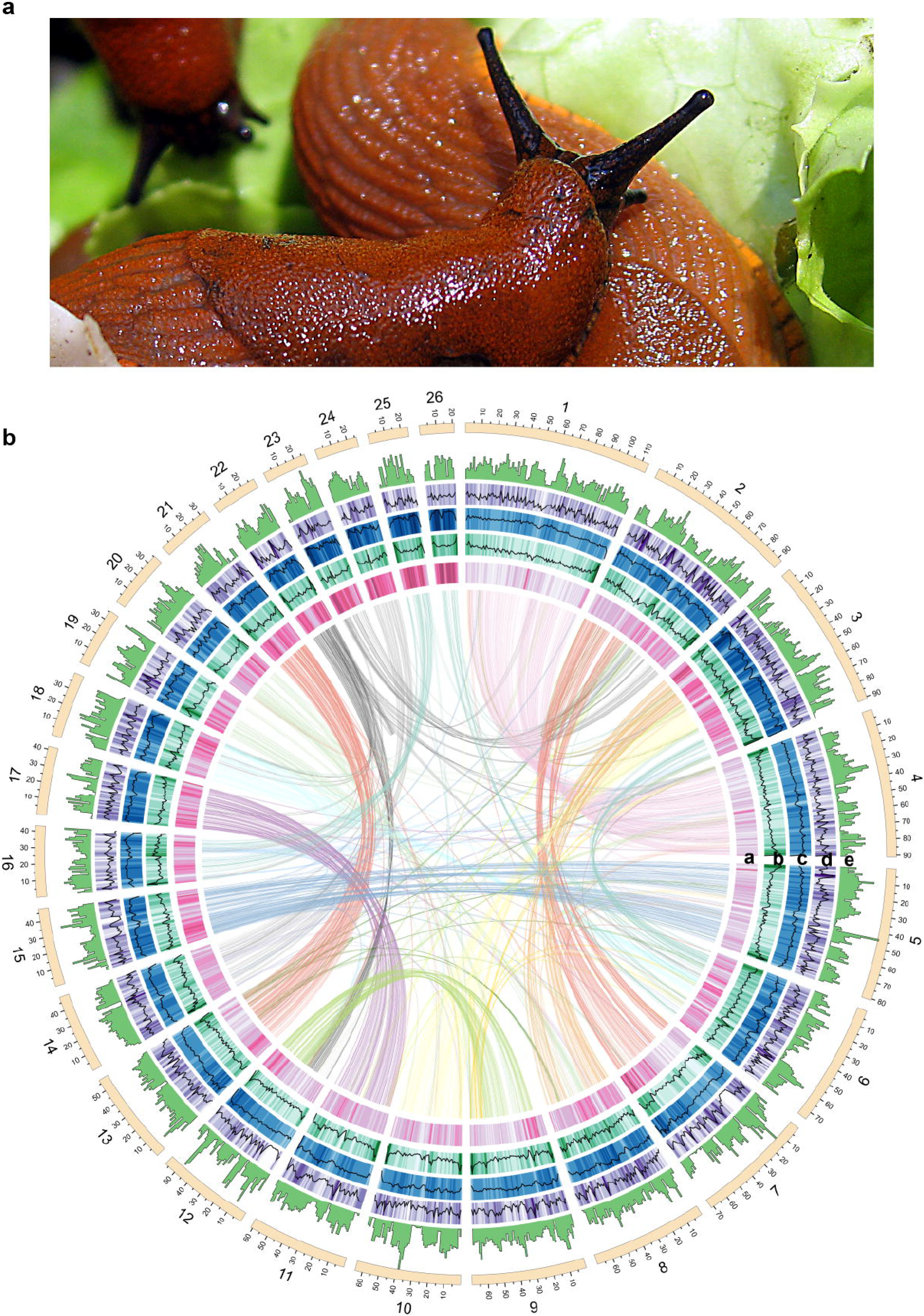
Genome features of *Arion vulgaris*. **(a)** Adult *A. vulgaris*. **(b)** General characteristics of the *A. vulgaris* genome. Tracks from inner to outside correspond to (a) GC content, (b) LTRs density, (c) TEs density, (d) genes density and (e) heterozygosity in sliding windows of 1 Mb across each of the 26 pseudochromosomes. Inner lines connect syntenic genes due to ancestral whole-genome duplication events. the graph was visualized by Circos v0.69-6 [36].

Total RNA was extracted from the ‘head’ part of the sample which includes tentacles, mantle, inner head and anterior visceral organs, and foot and sequenced on an Illumina NovaSeq platform with paired-end 150 bp. ~7 Gb reads were generated and further used for aiding genome annotation (**Table S1**).

### Genome feature estimation and assembly

The genome size and heterozygosity were estimated by GenomeScope v1.0.0 [30] using the quality-controlled paired-end Illumina sequence data and linked reads. The estimated genome size is 1.45 Gb with a high heterozygosity of 1.03% and repeat content of 52.98% in the genome **(Fig S1)**.

Long reads, generated with the Nanopore PromethION platform, were assembled into contigs using the wtdbg2 v2.2 assembler [31]. The contigs were subsequently polished by ntEdit v1.3.1 using Illumina short reads and linked reads [32]. The resulting contigs were then connected to super-scaffolds by 10X Genomics linked-read data using the Scaff10X v4.2 software. These processes yielded a draft *A. vulgaris* genome assembly with a total length of 1.54 Gb in 6,109 contigs, and with a contig N50 length of 4.49 Mb and a scaffold N50 length of 7.66 Mb **(Table S2)**.

Hi-C reads were used to generate a chromosome-level assembly of the genome. Hi-C reads were mapped to the draft assembly and processed using the hicstuff v2.2.2 pipeline [33] with the parameters --aligner bowtie2 --enzyme MboI --iterative --matfmt graal --quality-min 30 --size 0. instaGRAAL v0.1.2 was run on the resulting matrix and the draft assembly with parameters --level 5 --cycles 100 --coverage-std 1 --neighborhood 5 [34]. After building the interaction map of the final scaffolds with hicstuff, we noticed a relocation in chromosome 9 which may be due to a misassembly **(Fig S2a)**. In the subsequent analysis, we mapped all reads to the controversial region and specified two breakpoints (at site 21,000,000 and 27,600,000 respectively) based on the reads’ depth and gene distribution. We corrected the orders manually and reconnected sequences with 10 N’s at the new junction site **(Fig S2b)**. We obtained an assembly with a total length of 1.54 Gb (close to the estimated genome size, **Table S2**), a contig N50 of 8.6 Mb and a scaffold N50 of 63.3 Mb. 93.81% of the assembled genome was anchored onto 26 pseudochromosomes **(Table S2 and Fig 1b)**, which is consistent with the previous karyotype study [35, 36].

We assessed the quality of the genome assembly in three aspects: 1) more than 95.99% of the Illumina short reads could be mapped to the assembly by BWA-MEM v0.7.17-r1188 software with default parameters [37]; 2) a total of 886 (90.59%) conserved genes in BUSCO’s metazoan (odb9) benchmark set were present and complete in the genome (BUSCO v3.0.2) **(Table S2)**[38]; 3) the *k*-mer distribution showed a relatively collapsed assembly including mostly a single copy of the homozygous content and a partial representation of the heterozygous content, as is expected in a haploid assembly **(Fig S3)**, while comparing the clean reads spectrum of paired-end and linked reads libraries with the copy number in the assembly using KAT toolkit v2.4.2 [39]. These results all suggested a high-quality genomic resource of this initial genome assembly of *A. vulgaris* for subsequent analysis.

Heterozygosity was estimated by the following steps. First, the clean Illumina reads and linked reads were merged and mapped onto the assembly by BWA-MEM v 0.7.17-r1188 software with the default parameters [37]. Then, sequence alignment/map (SAM) format files were imported to SAMtools v1.9 for sorting and merging [40], and Picard v2.23.3 was used to mark duplicates [41]. Finally, single nucleotide polymorphisms (SNPs) calling was implemented in Genome Analysis Toolkit (GATK) v 4.1.6.0 using default parameters [42], and a number of filtering steps were performed to reduce the false positives, including: 1) remove SNPs with more than two alleles; 2) remove SNPs with quality score less than 30; 3) remove SNPs at or within 5bp from any InDels; 4) remove sites with extremely low (less than one-third average depth) or extremely high (more than three-fold average depth) coverage. We observed an average genome-wide heterozygosity rate of 1.55 per hundred base pair in *A. vulgaris*, which is about three times of the published most closely related species, *Achatina fulica* (Stylommatophora, heterozygosity rate: 0.47 per hundred base pair) [23], however, throughout all reported gastropod genomes (heterozygosity rate from 0.08 to 3.66 per hundred base pair) [5], its heterozygosity is at an intermediate level.

### Repeat prediction and expansions of transposable elements

Tandem Repeat Finder v4.09 (TRF) was used for tandem repeats identification with default parameters [43]. Transposable elements (TEs) were annotated using a combination of *ab initio* and homology-based approaches. First, repeat elements were identified *de novo* using RepeatModeler v2.0.1, the result database, along with the RepBase (RepBase-20170127) and Dfam (Dfam_Consensus-20170127) libraries with all species [44, 45], were used as a custom library for RepeatMasker v4.0.7 to identify repeats comprehensively [46]. Finally, we confirmed that the repeat sequences occupy approximately 75.09% (1.15 Gb) of the assembly **(Table S3)**, which is the highest value among all studied gastropod species [25]. We also found that species in Heterobranchia have higher repeat content than other gastropod groups (i.e., Caenogastropoda, Vetigastropoda, Neomphaliones, and Patellogastropoda) **(Fig S4)**. In all types of repetitive sequences, TEs account for 61.08% of the *A. vulgaris* assembly, among them, long interspersed elements (LINEs), DNA transposons (DNAs), short interspersed elements (SINE) account for 36.39%, 5.44%, 1.78% of the assembly respectively, Unknown TEs account for a high proportion (17.76%) of the genome (**Table S3**). The repeat divergence rate was measured by the percentage of substitutions in the corresponding regions between annotated repeats and consensus sequences in the RepBase database. We conducted a comprehensive comparison of the proportion of the different type of TEs and their divergence rate among *A. vulgaris* and other published Heterobranchia species (*Ac. fulica* [23], *Ac. immaculata* [22], *Biomphalaria glabrata* [47], *Radix auricularia* [48], *Aplysia californica*, and *Elysia chlorotica* [49]). Notably, the main class of TEs in *A. californica*, *E. chlorotica* and *R. auricularia* are DNA while in *A. vulgaris*, *Ac. fulica*, *Ac. immaculata*, and *B. glabrata*, LINEs are dominant (**Fig 2a**). Moreover, DNAs and LINEs showed a considerable amount with a low divergence rate in *A. vulgaris*, indicating a recent explosion of DNAs and LINEs in the *A. vulgaris* genome (**Fig 2b**). Previous reports have shown that TEs are powerful facilitators of rapid adaptation that generate “evolutionary potential” by introducing stress-induced changes in invasive species [50], and a recent genomic study of invasive land snails observed recent TE explosions in four invasive mollusk species (*Ac. fulica*, *Ac. immaculata*, *Pomacea canaliculate*, and *Crassostrea gigas*) while absent in other, noninvasive mollusks [22]. Significantly, the expansion of these two types of transposons in *A. vulgaris* is more obvious than the recently reported TEs expansion in two invasive snails *Ac. fulica* and *Ac. immaculata* **(Fig 2b)**. Therefore, the recent expansion of DNA and LINE TEs in *A. vulgaris* may also play important roles in promoting potential plasticity and stress resistance correlated with its invasiveness and competitiveness.

**Figure 2.**
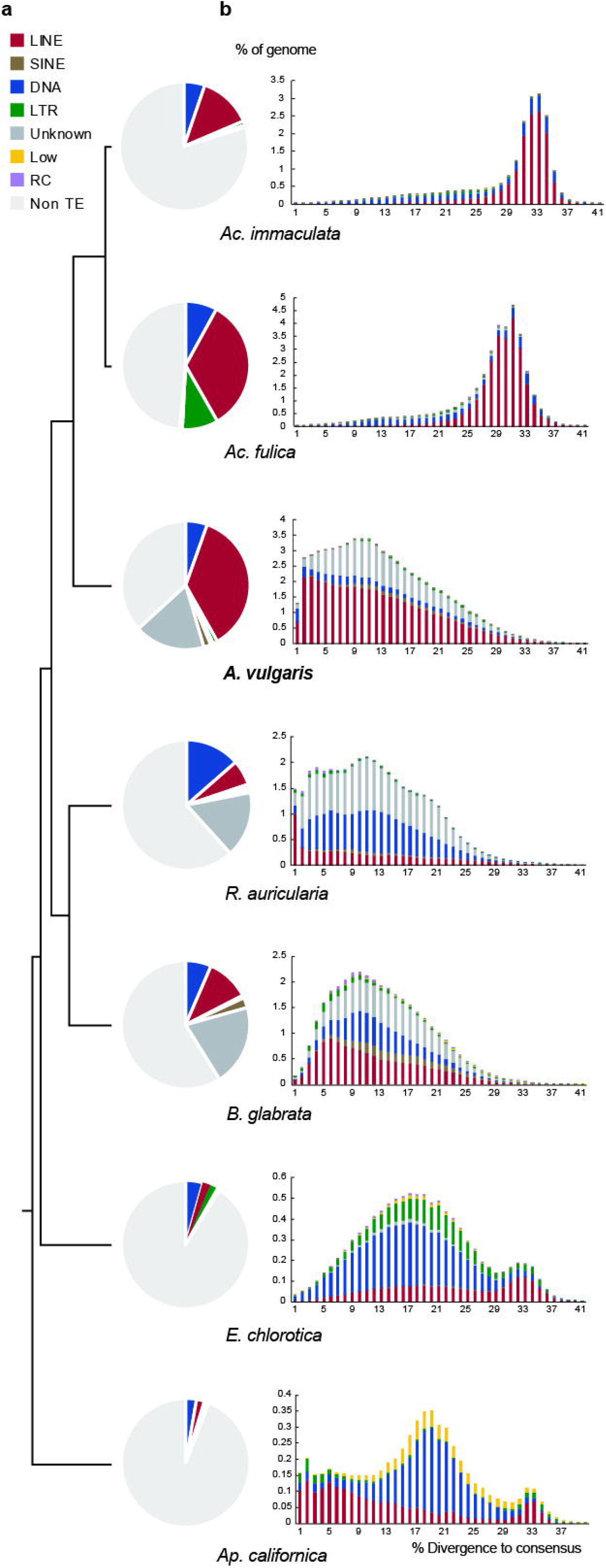
Transposable element (TE) landscape among published Heterobranchia species. **(a)** TE composition by class (indicating % of the genome corresponding to each class). **(b)** TE accumulation history. Cumulative amounts (% of genome size) are shown as a function of the percentage of divergence from the consensus. The percentage of divergence from consensus is used as a proxy for age. The older the invasion of the TE is, the more copies will have accumulated mutations, the higher percentage of divergence, right to the graph and vice versa.

### Gene prediction

Protein-coding genes were predicted using the following approaches: *ab initio* prediction, homology-based prediction, and transcriptome-based prediction. For *ab initio* prediction, RNA-seq reads were first aligned to the *A. vulgaris* genome sequence using STAR v2.7.2b [51], then the read alignment information was merged and used for Braker2 v2.1.5 gene prediction pipeline [52]. For homology-based prediction, protein sequences of six mollusk species (*Ac. fulica* [23], *B. glabrata* [47], *Ap. californica*, *E. chlorotica* [49], *P. canaliculata* [53], *Haliotis rufescens* [54]) were downloaded from NCBI and aligned against the assembled genome with MMseqs v 11.e1a1c [55]. These results were then combined into gene models separately with GeMoMa v1.3.1 using mapped RNA-seq data for splice site identification [56]. The resulting gene annotation sets were further filtered using the GeMoMa module GAF using default parameters. For the transcriptome-based prediction, RNA-seq data had been assembled in both *de novo* and genome-guided way with Trinity vr20140413p1, and the gene predictions were carried out with Program to Assemble Spliced Alignments (PASA) v2.0.2 [57, 58]. All gene annotations were joined with EvidenceModeller (EVM) v1.1.1 (**Table S4 and S5**) [59]. Moreover, partial genes and genes with a coding length of less than 150 bp were further filtered. The final gene set contains 32,518 predicted genes with an average length of 15,429 bp (**Table S5 and Fig S5**). The length distribution of transcripts, coding sequences, exons and introns, and the distribution of exon numbers per gene were comparable between *Ac. fulica, Ac. immaculata, B. glabrata* and *R. auricularia.* (**Fig S6**).

The predicted genes were annotated by aligning them to the eggNOG [60], SWISS-PROT [61], TrEMBL [61], KEGG [62], and InterPro [63] databases using BLAST v2.2.31 with a maximal e-value of 1e−5 [64] and by aligning to the Pfam [65] database using HMMer v3.0 [66]. Gene Ontology (GO) terms (Gene Ontology, RRID:SCR 002811) were assigned to the genes using the BLAST2GO v2.5 pipeline [67]. As a result, a total of 31,763 genes were annotated in one or more databases **(Table S6)**. A total of 11,262 and 9,642 genes were annotated in the GO and KEGG databases, respectively **(Table S6)**.

### Gene family clustering and phylogenetic analysis

We used protein sequences from all gastropods that have published genomes (except for *Achatina*, for which we only retained one species), covering five subclasses of gastropods, including six Heterobranchia: *Ac. fulica* [23], *Ac. immaculata* [22], *B. glabrata* [47], *R. auricularia* [48], *Ap. californica*, *E. chlorotica* [49], four Caenogastropoda: *P. canaliculata* [53], *Marisa cornuarietis* [53], *Lanistes nyassanus* [53], *Conus consors* [68], one Vetigastropoda: *H. rufescens* [54], one Neomphaliones: *Chrysomallon squamiferum* [69, 70] and one Patellogastropoda: *Lottia gigantea* [71], as well as the two bivalve species *Argopecten purpuratus* [72] and *Saccostrea glomerata* [73] were used as outgroups cluster gene families and construct phylogenetic tree. An all-against-all comparison was performed using BLASTP with an e-value cut-off of 1e-5 [74], and OrthoFinder v2.4.0 was used to cluster gene families [75]. We identified 30,636 *A. vulgaris* genes clustered into 13,333 families **(Table S7)**. Among these, 7,688 orthologous gene families were shared by five Panpulmonata species, whereas 1,126 gene families were exclusively shared by *A. vulgaris*, *Ac. fulica* and *Ac. immaculata* **(Fig 3a)**. There was a total of 2,763 genes unique to *A. vulgaris* **(Fig 3b and Table S7)**and 2,629 (95.2%) of which have known InterPro domains. We found these *A. vulgaris* specific genes were significantly enriched in functional categories including response to different stimuli, e.g. chemical stimulus (GO:0070887), abiotic stimulus (GO:0009628), external stimulus (GO:0009605), positive regulation of response to stimulus (GO: 0048584), signal transduction (GO:0023052, GO:0009966, GO:0007165), neurons generation (GO:0048699, GO:0048699) and differentiation (GO:0030182), muscle structure development (GO:0061061) and embryo development (GO:0009792) **(Table S8)**, which might benefit to their high fertility, ecological tolerance and plasticity. Some of the species-specific genes are related to the immune system, but compared to all other genes, these are not significantly enriched (P>0.05).

**Figure 3.**
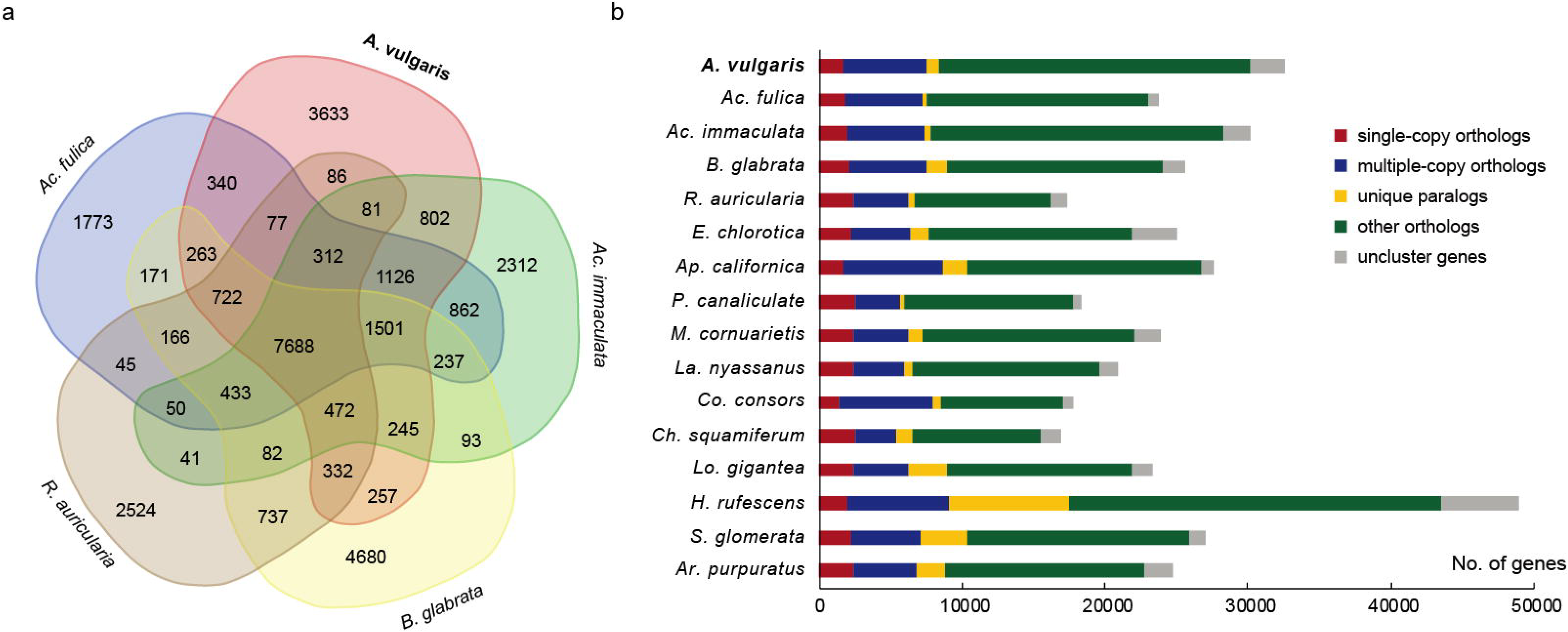
Gene family analyses of the *Arion vulgaris* genome. **(a)** Venn diagram showing the number of shared and unique gene families among five Panpulmonata genomes. **(b)** Gene clusters of 16 molluscan species, bars represent the number of genes in different categories for each species.

The 1,237 gene families with only one copy from each of 16 species were selected as single-copy genes and were used for subsequent analysis. We aligned these protein sequences using MUSCLE v3.8.1551 [75] and constructed the phylogenetic tree using RAxML v8.2.8 with the GTR+ Γ model assigned to each partition [76]. The divergence time of each species was computed using the MCMCTREE program implemented in the Phylogenetic Analysis by Maximum Likelihood (PAML v4.8) package [77]. For calibration we used the soft bounds of Euopisthobranchia – Panpulmonata (divergence time between 190 MY and 270 MY) [78], the fossil of Sublitoidea (418 MY) constraints on the node of Heterobranchia and Caenogastropoda [79], and the fossil of *Fordilla troyensis* (530 MY) for the root [80, 81]. As a result, *A. vulgaris* diverged from a stylommatophoran branch with two land snails *Ac. fulica* and *Ac. immaculata* ~126 MY (slightly earlier than the estimation based on mitochondria genome data, 132 MY [82]), and the combined clade separated from a common ancestor with the hygrophilan *B. glabrata*, and *R. auricularia* around 199 MY **(Fig 4)**.

**Figure 4.**
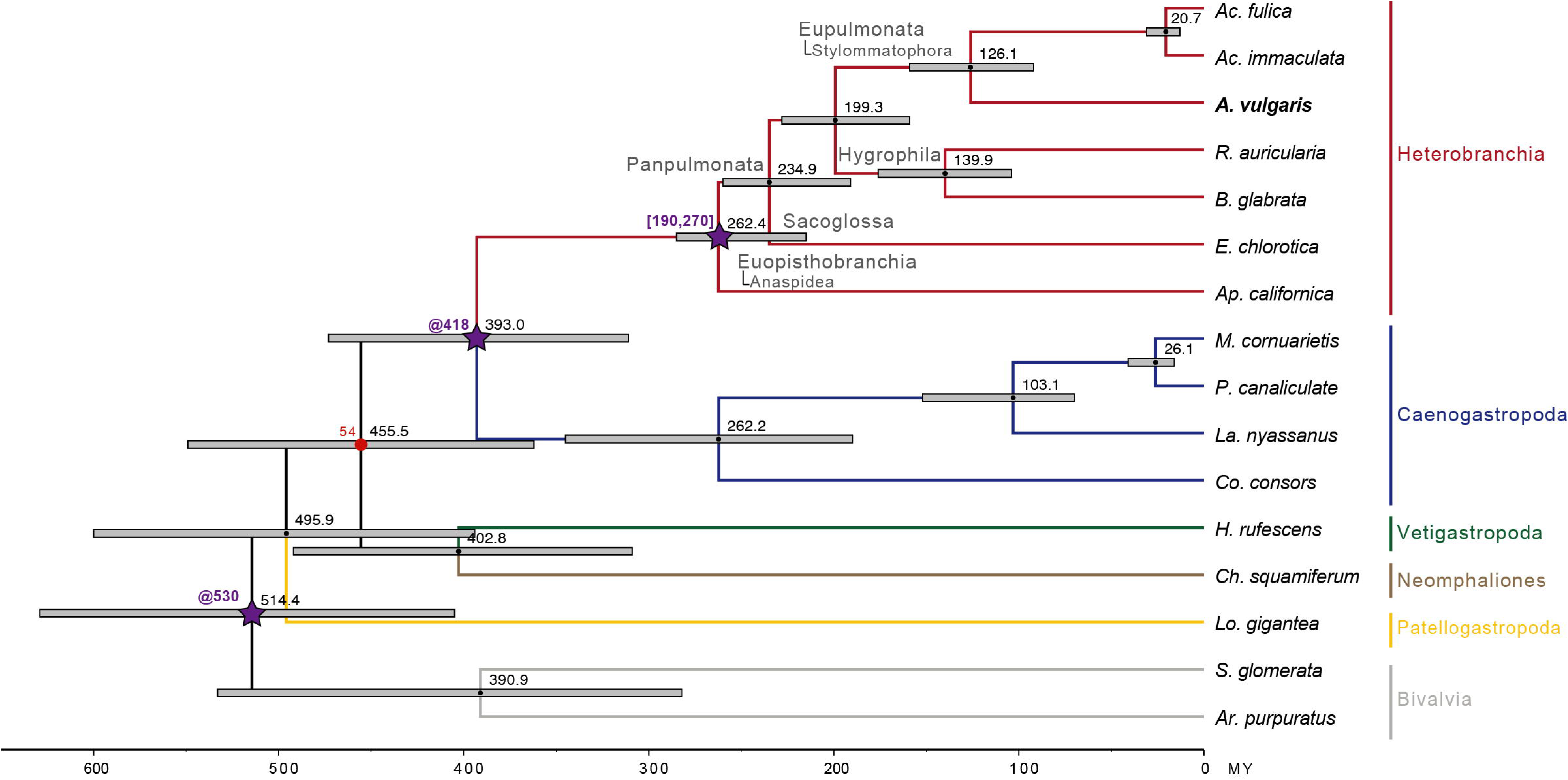
Dated Maximum-likelihood phylogenetic tree among gastropod species with whole-genome data available. The numbers on the nodes represent the divergence times from the present (million years ago, MY). Blue lines indicate the 95% confidence interval on the estimated node depths. Purple stars indicate that nodes are constrained with fossil calibration or secondary calibration data. All nodes have a bootstrap support values of 100 except the red node with a low bootstrap of 54.

### Identification of whole-genome duplication event

Multiple sequence alignment was conducted in *A. vulgaris* and also in *Ac. fulica* and *A. immaculata*. The protein sequences were first searched against themselves using BLASTP v2.9.0 [74] and then subjected to MCScan v0.8 [83] to determine syntenic blocks. The result of collinearity shows that most of the chromosomes can be found the corresponding one in the *A. vulgaris* genome and a one-to-one corresponding relationship in the comparison of *A. vulgaris* and *A. immaculata* **(Fig 1 and Fig S6)**. We then used add_kaks_to_synteny.pl, the downstream analysis of MCscanX v0.8 [84] to calculate the synonymous substitutions of each pair of genes in the syntenic blocks. Frequency distributions are shown of values of synonymous substitutions (*K*S) for paralogous and orthologous genes in comparisons of *A. vulgaris (A. vu)*, *Ac. fulica (A. fu)*, and *Ac. immaculata (A. im)*. Peaks in the *A. vu*–*A. vu, A. fu-A. fu, A. im-A. im* comparisons represent the whole-genome duplication (WGD). Assuming that the mutation rate of Mollusca is 1.645×10^−9^ per site per year [85], the peaks of *A. vu*–*A. vu, A. fu-A. fu* are at *K*s = 1.16 and *K*s = 1.26, respectively, the estimated WGD event time is around 70-77 MY **(Fig 5)**. The peak of A. im-A. im, however, is at Ks = 0.65 (~40 MY), which is earlier than expected **(Fig 5)**. A previous study has proved the occurrence of a WGD event shared by *Ac. immaculata* and *Ac. fulica* around 70 MY [22], and another earlier study speculated the WGD event somewhere at the base of Stylommatophora by comparison of chromosome numbers among closely related mollusks [86]. The reason why *Ac. immaculata* does not have the same peak as *Ac. fulica* and *A. vulgaris* remains to be further discussed, however, our results contribute to another genomic evidence for the occurrence of the WGD event in Stylommatophora.

**Figure 5.**
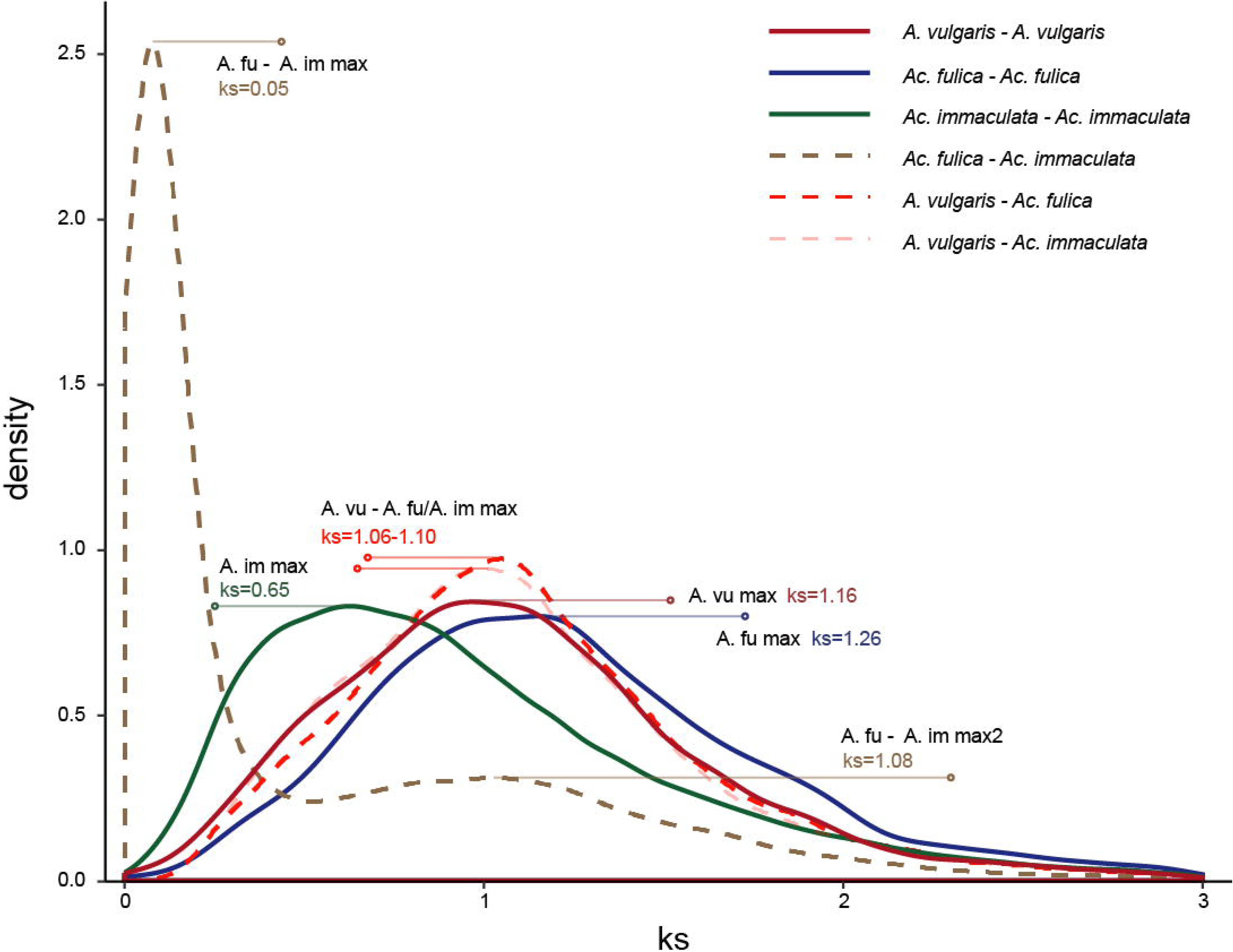
Whole-genome duplication shared by *Arion vulgaris*, *Achatina fulica* and *Ac. immaculata*. Frequency distributions are shown of values of synonymous substitutions (Ks) for paralogous and orthologous genes in comparisons of *A. vulgaris (A.vu)*, *Ac. fulica (A.fu)* and *Ac. immaculata (A.im)*.

### Conclusions

We generated a chromosome-level assembly for the Spanish slug *A. vulgaris* using an integrated sequencing strategy combining 10X genomics, Nanopore, Illumina and Hi-C technologies. We predicted 32,518 genes and 97% of the genes were functionally annotated. We found recent DNA and LINE TEs were expanded in the *A. vulgaris* genome. The TE explosion, enrichment of species-specific genes in the function of response to stimuli, nervous system and reproduction might relate to its recent invasiveness. The high-quality genomic data provided here are valuable genetic resources for further *A. vulgaris* and molluscan studies.

## Availability of supporting data

The *A. vulgaris* genome project was deposited at the National Center for Biotechnology Information (NCBI) under BioProject number PRJNA680311.

## Additional Files

Fig S1. Estimation of genome size of *Arion vulgari*s based on the distribution of 21-mer frequency in the combination of short reads and linked reads.

Fig S2. Hi-C chromosome contact map.

Fig S3. Read *k-mer* frequency versus assembly copy number stacked histograms for final *Arion vulgaris* assembly.

Fig S4. Repetitive content in published gastropod genomes.

Fig S5. Comparison of mRNA length (a), CDS length (b), exon length (c), exon number per gene (d), and intron length (e), between *Arion vulgaris*, *Achatina fulica, Ac. immaculata, Biomphalaria glabrata,* and *Radix auricularia.*

Fig S6. A one to one corresponding relationship in the comparison of *Arion vulgaris* and *Achatina immaculata* genomes.

Table S1. Sequencing data generated for *Arion vulgaris* genome assembly and annotation.

Table S2. Summary of the assembly of *Arion vulgaris*.

Table S3. Identified repeat classes in the *Arion vulgaris* genome.

Table S4: Evidence weight file used for *Arion vulgaris* gene prediction.

Table S5: Statistics of predicted protein-coding genes in the *Arion vulgaris* genome. Table S6: Functional annotation of the predicted genes models.

Table S7: Summary of gene family clustering of 16 mollusk species.

Table S8: GO enrichment analysis of *Arion vulgaris* species-specific and unassigned genes.

## Abbreviations

BLAST: Basic Local Alignment Search Tool
BUSCO: Benchmarking Universal Single-Copy Orthologs
BWA: Burrows-Wheeler Aligner
CDS: coding DNA sequence
CTAB: Cetyltrimethylammonium-bromide
DAISIE: Delivering Alien invasive species inventoried for Europe
DNAs: DNA transposons
EVM: EvidenceModeller
GATK: Genome Analysis Toolkit
GO: gene ontology
KAT: the K-mer Analysis Toolkit
KEGG: Kyoto Encyclopedia of Genes and Genomes
LINEs: Long Interspersed Nuclear Elements
LTR: Long Terminal Repeat
MY: million years
NCBI: National Center for Biotechnology Information
PASA: Program to Assemble Spliced Alignments
RC: Rolling Circle
SAM: Sequence Alignment Map
SINEs: Short Interspersed Nuclear Elements
SNPs: single nucleotide polymorphisms
TEs: transposable elements
TrEMBL: Translation of European Molecular Biology Laboratory
TRF: Tandem Repeat Finder
WGD: whole-genome duplication

## Consent for publication

Not applicable.

## Ethics Statement

No specific permits were required for the described field studies, no specific permissions were required for these locations/activities, and the field studies did not involve endangered or protected species.

## Author contributions

Z.C. and M.S. conceived the study. Z.C., Ö.D., N.D., A.G., performed genome assembly. Z.C. performed genome annotation and evolution analysis. Z.C. and M.S. wrote the manuscript. All authors contributed to editing the manuscript.

## Acknowledgements

We thank Gert Wörheide and Michael Eitel (Department of Earth and Environmental Sciences, Ludwig-Maximilians-Universität München, LMU) for organizing the IGNITE Comparative Genomics of Non-Model Invertebrates program; Denis Tagu (INRAE), Fabrice Legeai (INRAE, INRIA), Claire Lemaitre (INRIA), Grace McCormack and Kenneth Sandoval (National University of Ireland) for the guidance and advice on genome assembly during secondments; Ramon Rivera (LMU) for software installation and maintenance; Juan Moles (ZSM) for polishing the article, and other ZSM mollusk teammates as well as colleagues from IGNITE for valuable feedback and comments. The bioinformatics analyses were performed at Leibniz-Rechenzentrum der Bayerischen Akademie der Wissenschaften, TUBITAK ULAKBIM, High Performance and Grid Computing Center (TRUBA resources) and GenOuest IRISA-INRIA, France.

