## Supplement Figures and Tables for "The *de novo* genome of the “Spanish” slug *Arion vulgaris* Moquin-Tandon, 1855 (Gastropoda: Panpulmonata): massive expansion of transposable elements in a major pest species"

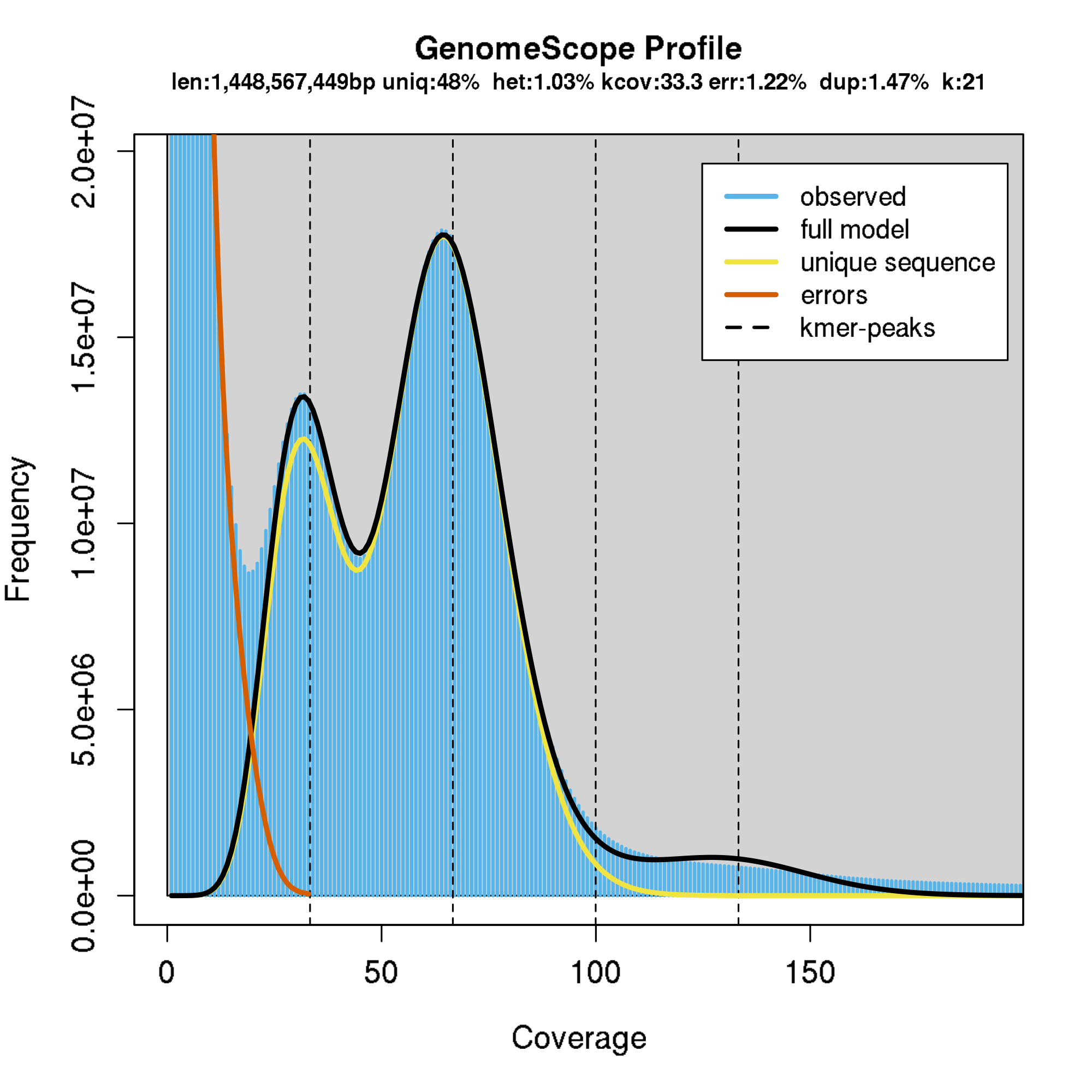

**Fig S1. Estimation of genome size of *Arion vulgari*s based on the distribution of 21-mer frequency in the combination of short reads and linked reads.** Two peaks indicate high heterozygosity.

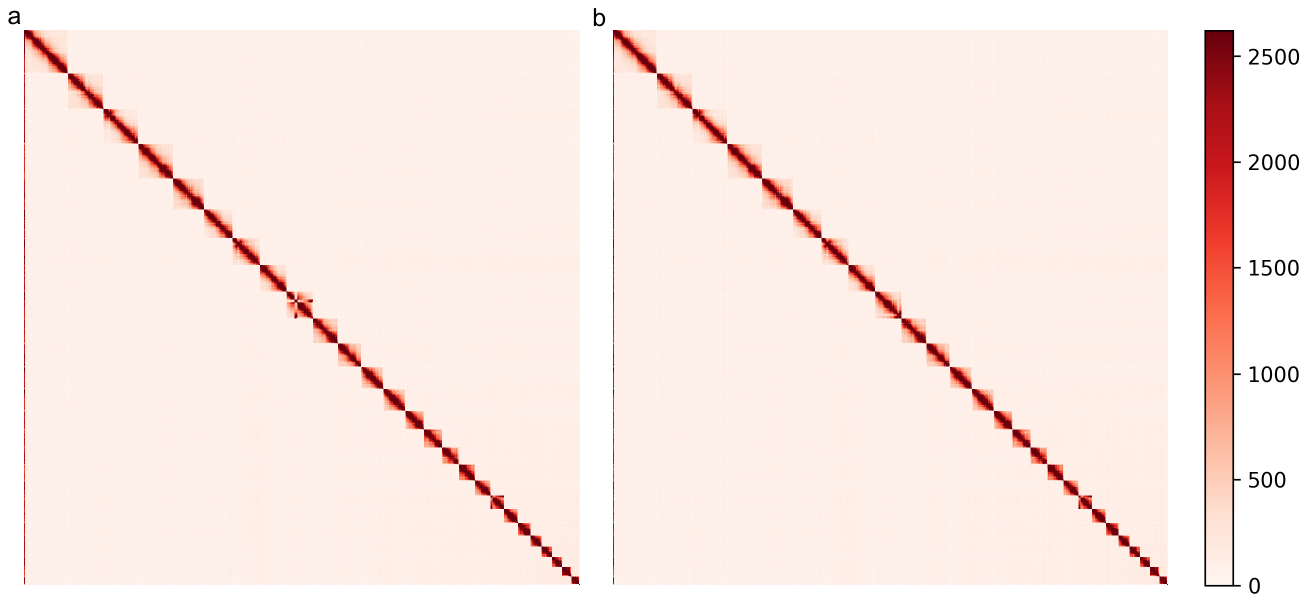

**Fig S2. Hi-C chromosome contact map.** Each block represents a Hi-C contact between two genomic loci within a 1 Mb window. Darker color of a block indicates higher contact intensity. a) Hi-C contact map shows incongruous in chromosome 9 which might cause by the misassembly. b) Hi-C contact map after manually corrected the assembly.

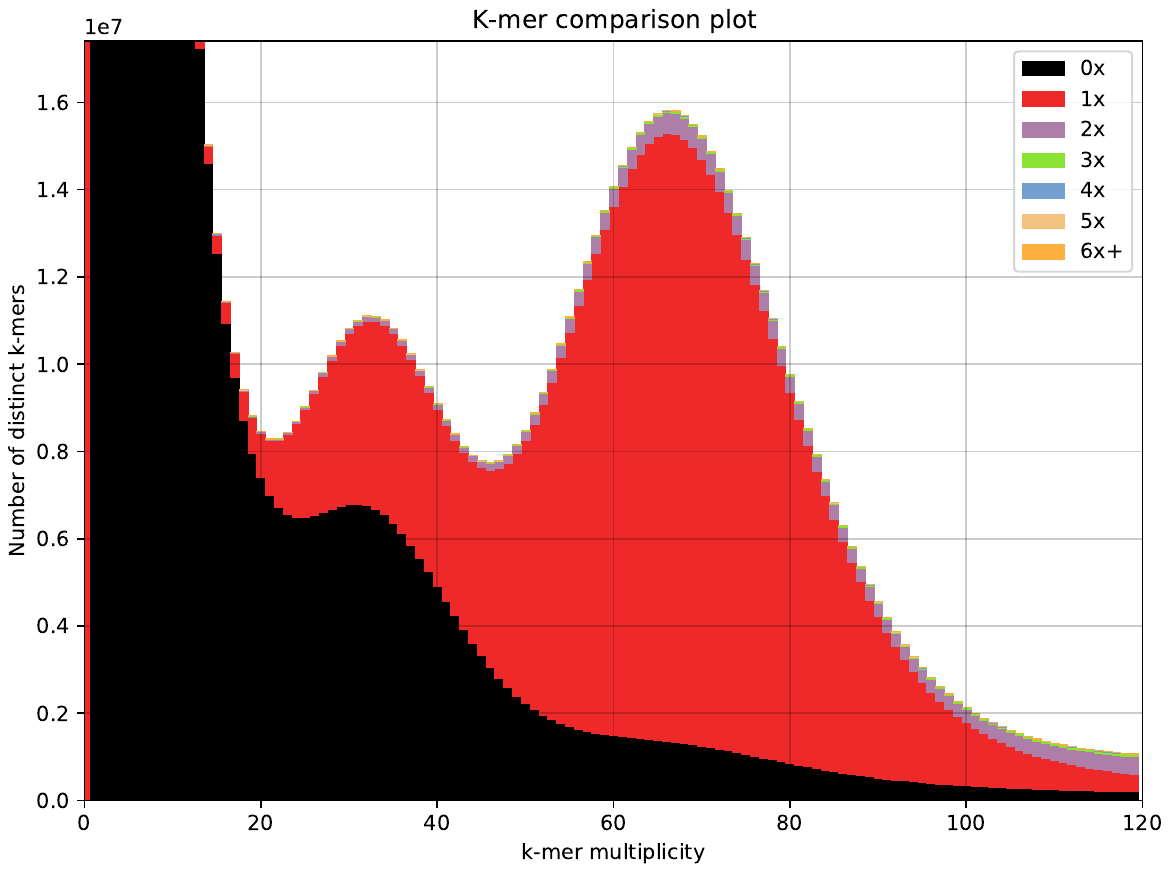

**Fig S3. Read *k-mer* frequency versus assembly copy number stacked histograms for final *Arion vulgaris* assembly.** Read content in black is absent from the assembly, red occurs once, purple twice, etc.

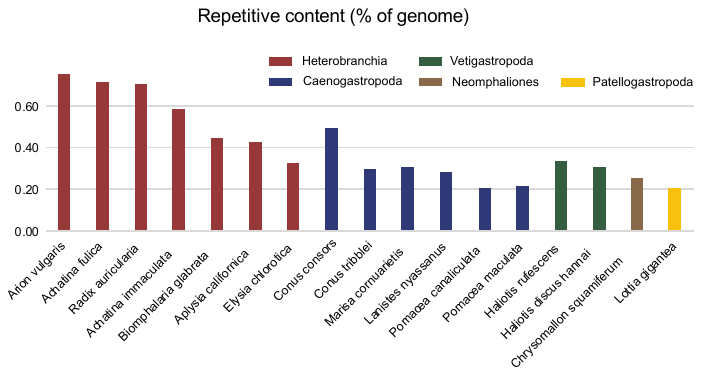

**Fig S4. Repetitive content in published gastropod genomes.**

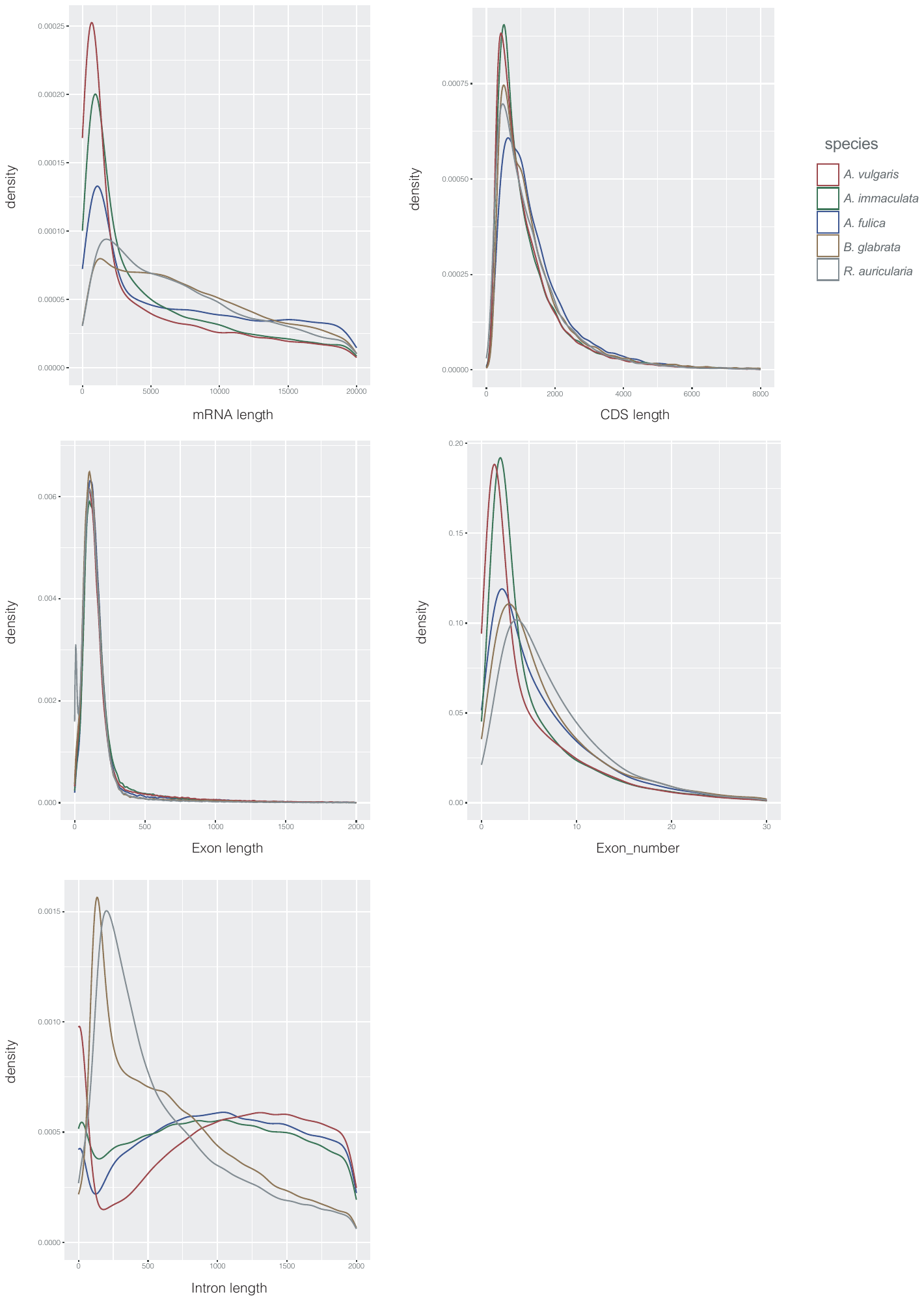

**Fig S5. Comparison of mRNA length (a), CDS length (b), exon length (c), exon number per gene (d), and intron length (e), between *Arion vulgaris*, *Achatina fulica, Ac. immaculata, Biomphalaria glabrata,* and *Radix auricularia.***

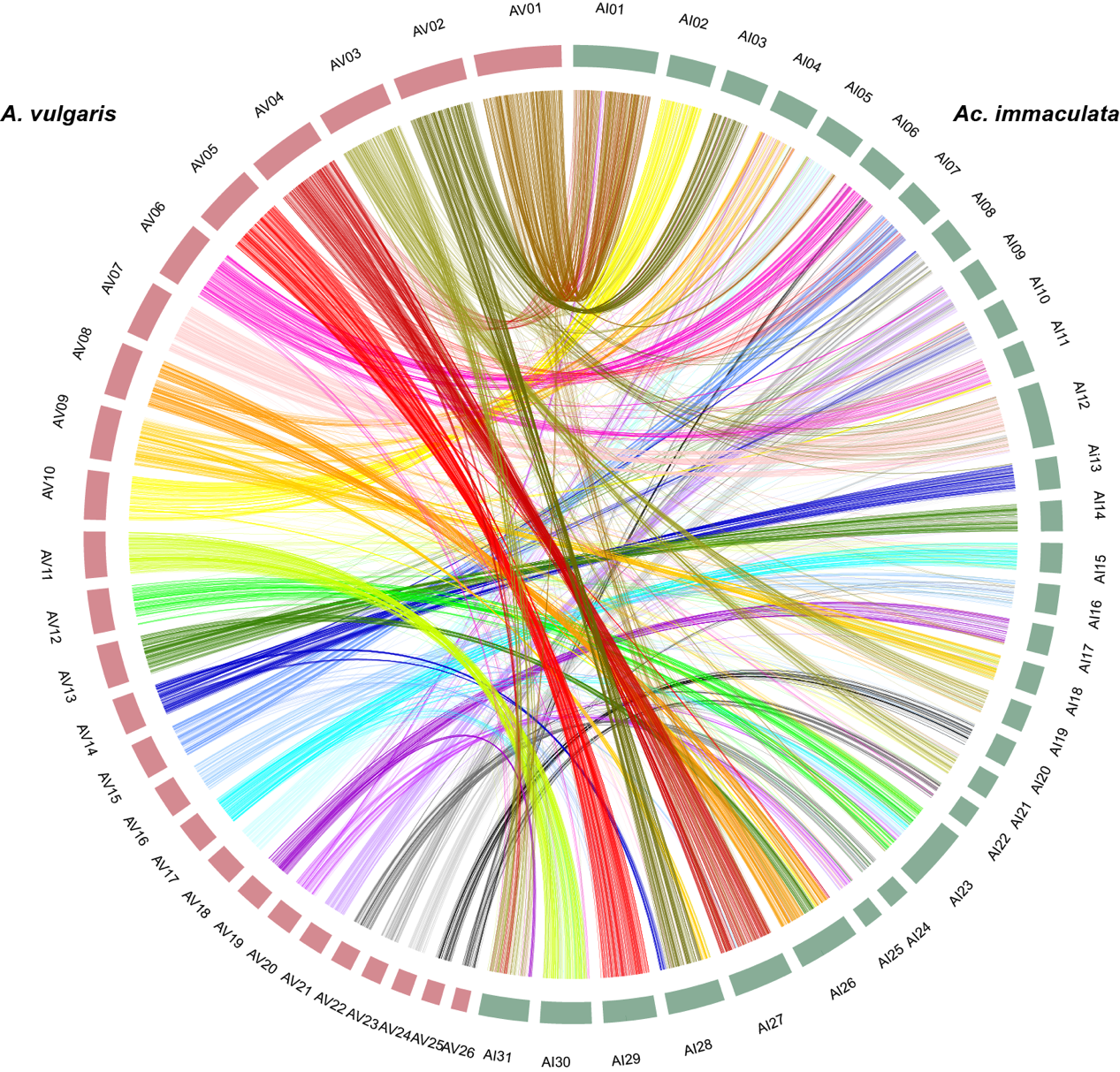

**Fig S6. A one to one corresponding relationship in the comparison of *Arion vulgaris* and *Achatina immaculata* genomes*.***

**Table S1. Sequencing data generated for *Arion vulgaris* genome assembly and annotation.**

| Library type | Platform | Total bases  (Gb) | Application |
| --- | --- | --- | --- |
| Short reads | HiSeq X Ten | 56.83 | Genome survey and genomic base correction |
| Linked reads | HiSeq X Ten | 138.21 | Genome survey, genomic base correction and genome assembly |
| Nanopore | Nanopore PromethION | 74.94 | Genome assembly |
| HiC | HiSeq X Ten | 135.28 | Chromosome construction |
| mRNA-Seq | NovaSeq 6000 | 6.75 | Genome annotation |

**Table S2. Summary of the assembly of *Arion vulgaris*.**

| Characteristic | wtdbg2 | wtdbg2-ntEdit-Scaff10X | instaGRAAL |
| --- | --- | --- | --- |
|  | Nanopore reads | Nanopore reads, short reads, linked reads | Nanopore reads, short reads, linked reads, HiC reads |
| Total Contig length (Gb) | 1.535 (6,109) | 1.543 (6,109) | 1.543 (8,243) |
| Contig N50 (Mb) | 4.466 (93) | 4.488 (93) | 8.603 (49) |
| Longest Contig (Mb) | 23.976 | 24.090 | 39.732 |
| Total scaffold length (Gb) | - | 1.543 (5,786) | 1.543 (7,920) |
| Scaffold N50(Mb) | - | 7.660 (55) | 64.342 (10) |
| Longest Scaffold (Mb) | - | 34.170 | 114.239 |
| N's per 100 kbp (bp) | - | 2.09 | 2.09 |
| Complete BUSCOs | 0.84 | 0.90 | 0.91 |
| Complete and single-copy BUSCOs | 0.82 | 0.84 | 0.85 |
| Complete and duplicated BUSCOs | 0.02 | 0.06 | 0.06 |
| Fragmented BUSCOs (F) | 0.03 | 0.02 | 0.02 |
| Missing BUSCOs (M) | 0.13 | 0.08 | 0.07 |

Note: The number in brackets indicates the number of corresponding scaffolds and contigs.

**Table S3. Identified repeat classes in the *Arion vulgaris* genome.**

| Repeat Type | | Repeat Size (bp) | %of Genome |
| --- | --- | --- | --- |
| **TEs** | | 941,228,221 | 61.08 |
|  | LINE | 561,000,000 | 36.39 |
|  | DNA | 83,828,750 | 5.44 |
|  | SINE | 27,404,787 | 1.78 |
|  | LTR | 17,654,456 | 1.15 |
|  | RC | 10,002,034 | 0.65 |
|  | Satellite | 8,990,082 | 0.58 |
|  | low | 864,980 | 0.06 |
| Unknown | | 274,000,000 | 17.76 |
| **Tandem Repeats** | | 326,134,855 | 21.14 |
| **Total** | | 1,158,337,369 | 75.09 |

**Table S4: Evidence weight file used for *Arion vulgaris* gene prediction.**

| Item | | weight |
| --- | --- | --- |
| protein | |  |
|  | Achatina fulica | 4 |
|  | Biomphalaria globrata | 2 |
|  | Elysia chlorotica | 2 |
|  | Haliotis rufescens | 2 |
|  | Pomacea canaliculata | 2 |
|  | Aplysia california | 2 |
| abinitio prediction | | 1 |
| transcript | | 6 |

Table S5: Statistics of predicted protein-coding genes in the *Arion vulgaris* genome.

| Gene set | | Number | Averge transcript  length(bp) | Averge CDs  length(bp) | Average exon  per gene | Average exon  length(bp) | Averge intron  length(bp) |
| --- | --- | --- | --- | --- | --- | --- | --- |
| homolog prediction | |  |  |  |  |  |  |
|  | *Achatina fulica* | 22,718 | 10,568.96 | 1,202.44 | 4.60 | 261.53 | 2,603.49 |
|  | *Aplysia california* | 12,046 | 15,833.02 | 1,412.34 | 7.39 | 191.22 | 2,258.25 |
|  | *Biomphalaria globrata* | 16,264 | 15,628.49 | 1,325.09 | 7.06 | 187.81 | 2,362.09 |
|  | *Elysia chlorotica* | 13,069 | 13,604.99 | 1,204.99 | 6.42 | 187.82 | 2,289.63 |
|  | *Haliotis rufescens* | 21,393 | 5,791.92 | 863.26 | 2.82 | 305.68 | 2,701.96 |
|  | *Pomacea canaliculata* | 10,993 | 21,930.11 | 1,520.19 | 9.41 | 161.60 | 2,427.71 |
| *De novo* prediction | | 44,246 | 12,668.01 | 1,146.02 | 5.08 | 225.44 | 2,821.68 |
| transcriptome prediction | | 161,478 | 62,36.37 | 910.84 | 2.14 | 424.68 | 4,652.03 |
| final gene set | | 32,518 | 15,429.25 | 1,291.68 | 5.70 | 226.43 | 3,005.06 |

**Table S6: Functional annotation of the predicted genes models.**

|  | Number | Percent (%) |
| --- | --- | --- |
| eggNOG-mapper | 16,791 | 0.51636017 |
| KEGG | 9,642 | 0.296512701 |
| InterPro | 26,520 | 0.815548312 |
| SwissProt | 31,728 | 0.975705763 |
| TrEMBL | 31,429 | 0.966510856 |
| Total | **31,763** | 0.97678209 |

**Table S7:** Summary of gene family clustering of 16 mollusk species.

| species | total genes | genes in family | unassigned genes | family number | unique family | genes in unique family | genes  per family |
| --- | --- | --- | --- | --- | --- | --- | --- |
| *A. vulgaris* | 32,518 | 30,636 | 1,882 | 13,333 | 253 | 881 | 2.30 |
| *A. fulica* | 23,726 | 23,073 | 653 | 12,390 | 79 | 310 | 1.86 |
| *A. immaculata* | 30,194 | 28,633 | 1,561 | 12,513 | 124 | 361 | 2.29 |
| *B. globrata* | 25,550 | 24,093 | 1,457 | 12,444 | 366 | 1,450 | 1.94 |
| *R. auricularia* | 17,338 | 16,237 | 1,101 | 10,310 | 110 | 367 | 1.57 |
| *A. california* | 27,576 | 26,761 | 815 | 12,191 | 396 | 1,601 | 2.20 |
| *E. chlorotica* | 24,980 | 21,936 | 3,044 | 12,980 | 312 | 1,255 | 1.69 |
| *P. canaliculate* | 18,273 | 17,728 | 545 | 12,042 | 47 | 159 | 1.47 |
| *M. cornuarietis* | 23,827 | 21,988 | 1,839 | 13,181 | 201 | 1,005 | 1.67 |
| *L. nyassanus* | 20,938 | 19,651 | 1,287 | 12,475 | 138 | 555 | 1.58 |
| *C. consors* | 17,715 | 17,023 | 692 | 7,654 | 224 | 611 | 2.22 |
| *C. squamiferum* | 16,917 | 15,498 | 1,419 | 9,904 | 252 | 1,047 | 1.56 |
| *L. gigantea* | 23,340 | 21,940 | 1,400 | 12,165 | 408 | 2,794 | 1.80 |
| *H. rufescens* | 48,956 | 44,060 | 4,896 | 14,580 | 1,622 | 8,303 | 3.02 |
| *S. glomerata* | 26,956 | 26,327 | 629 | 11,516 | 619 | 3,375 | 2.29 |
| *A. purpuratus* | 24,705 | 22,797 | 1,908 | 12,568 | 501 | 2,069 | 1.81 |

**Table S8: GO enrichment analysis of *Arion vulgaris* species specific and unassigned genes**.

For each GO subcategory, a 2 × 2 contingency table was constructed by recording the numbers of genes included or not included in a category of ‘genome background’ genes and expanded genes. Two-tailed Fisher’s exact test was used to calculate statistical significance.

| GO ID | Description | P value |
| --- | --- | --- |
| GO:0070887 | cellular response to chemical stimulus | 9.01E-03 |
| GO:0016192 | vesicle-mediated transport | 5.54E-03 |
| GO:0044248 | cellular catabolic process | 4.63E-03 |
| GO:0034654 | nucleobase-containing compound biosynthetic process | 4.02E-03 |
| GO:0006464 | cellular protein modification process | 3.92E-03 |
| GO:0036211 | protein modification process | 3.92E-03 |
| GO:0050790 | regulation of catalytic activity | 3.62E-03 |
| GO:0009628 | response to abiotic stimulus | 3.50E-03 |
| GO:0022402 | cell cycle process | 3.48E-03 |
| GO:0030182 | neuron differentiation | 3.29E-03 |
| GO:0009792 | embryo development ending in birth or egg hatching | 2.53E-03 |
| GO:0044271 | cellular nitrogen compound biosynthetic process | 2.47E-03 |
| GO:0032268 | regulation of cellular protein metabolic process | 2.28E-03 |
| GO:0048729 | tissue morphogenesis | 1.99E-03 |
| GO:0034645 | cellular macromolecule biosynthetic process | 1.92E-03 |
| GO:0015031 | protein transport | 1.56E-03 |
| GO:0010605 | negative regulation of macromolecule metabolic process | 1.48E-03 |
| GO:0045184 | establishment of protein localization | 1.28E-03 |
| GO:0006629 | lipid metabolic process | 1.22E-03 |
| GO:0031324 | negative regulation of cellular metabolic process | 1.10E-03 |
| GO:0009059 | macromolecule biosynthetic process | 8.97E-04 |
| GO:0045595 | regulation of cell differentiation | 8.82E-04 |
| GO:0003008 | system process | 8.29E-04 |
| GO:0032989 | cellular component morphogenesis | 8.28E-04 |
| GO:0065003 | macromolecular complex assembly | 7.01E-04 |
| GO:0043933 | macromolecular complex subunit organization | 5.34E-04 |
| GO:0023056 | positive regulation of signaling | 5.33E-04 |
| GO:0009056 | catabolic process | 5.29E-04 |
| GO:0010647 | positive regulation of cell communication | 5.26E-04 |
| GO:0019438 | aromatic compound biosynthetic process | 4.67E-04 |
| GO:0043412 | macromolecule modification | 3.43E-04 |
| GO:0007049 | cell cycle | 3.27E-04 |
| GO:0051704 | multi-organism process | 2.90E-04 |
| GO:0016070 | RNA metabolic process | 2.89E-04 |
| GO:0010033 | response to organic substance | 2.64E-04 |
| GO:0051246 | regulation of protein metabolic process | 2.38E-04 |
| GO:0010467 | gene expression | 2.14E-04 |
| GO:1901575 | organic substance catabolic process | 1.87E-04 |
| GO:0060429 | epithelium development | 1.80E-04 |
| GO:0033036 | macromolecule localization | 1.80E-04 |
| GO:0018130 | heterocycle biosynthetic process | 1.58E-04 |
| GO:0045893 | positive regulation of transcription, DNA-templated | 1.35E-04 |
| GO:0009605 | response to external stimulus | 1.25E-04 |
| GO:1901362 | organic cyclic compound biosynthetic process | 1.18E-04 |
| GO:1902680 | positive regulation of RNA biosynthetic process | 8.87E-05 |
| GO:1903508 | positive regulation of nucleic acid-templated transcription | 8.87E-05 |
| GO:0071705 | nitrogen compound transport | 8.79E-05 |
| GO:0046907 | intracellular transport | 7.46E-05 |
| GO:0035295 | tube development | 6.50E-05 |
| GO:0061061 | muscle structure development | 4.95E-05 |
| GO:0008104 | protein localization | 4.62E-05 |
| GO:0009892 | negative regulation of metabolic process | 4.27E-05 |
| GO:0010557 | positive regulation of macromolecule biosynthetic process | 4.19E-05 |
| GO:0051649 | establishment of localization in cell | 4.08E-05 |
| GO:0071702 | organic substance transport | 3.15E-05 |
| GO:0044267 | cellular protein metabolic process | 2.56E-05 |
| GO:0048699 | generation of neurons | 1.93E-05 |
| GO:0051049 | regulation of transport | 1.71E-05 |
| GO:0050793 | regulation of developmental process | 1.58E-05 |
| GO:0048584 | positive regulation of response to stimulus | 1.58E-05 |
| GO:0009790 | embryo development | 1.09E-05 |
| GO:0051641 | cellular localization | 1.09E-05 |
| GO:0051254 | positive regulation of RNA metabolic process | 1.01E-05 |
| GO:0051128 | regulation of cellular component organization | 8.86E-06 |
| GO:0045935 | positive regulation of nucleobase-containing compound metabolic process | 5.20E-06 |
| GO:0009966 | regulation of signal transduction | 4.51E-06 |
| GO:0009887 | animal organ morphogenesis | 4.16E-06 |
| GO:0031328 | positive regulation of cellular biosynthetic process | 3.35E-06 |
| GO:0009891 | positive regulation of biosynthetic process | 3.04E-06 |
| GO:0033043 | regulation of organelle organization | 2.88E-06 |
| GO:0019538 | protein metabolic process | 1.83E-06 |
| GO:0090304 | nucleic acid metabolic process | 1.39E-06 |
| GO:0051239 | regulation of multicellular organismal process | 1.34E-06 |
| GO:0007165 | signal transduction | 1.28E-06 |
| GO:0022008 | neurogenesis | 1.04E-06 |
| GO:0006357 | regulation of transcription from RNA polymerase II promoter | 9.34E-07 |
| GO:0009888 | tissue development | 4.28E-07 |
| GO:0010646 | regulation of cell communication | 3.28E-07 |
| GO:0065009 | regulation of molecular function | 2.93E-07 |
| GO:0042592 | homeostatic process | 2.88E-07 |
| GO:0044249 | cellular biosynthetic process | 2.48E-07 |
| GO:0048468 | cell development | 2.10E-07 |
| GO:0006139 | nucleobase-containing compound metabolic process | 1.17E-07 |
| GO:0023051 | regulation of signaling | 1.07E-07 |
| GO:1901576 | organic substance biosynthetic process | 1.00E-07 |
| GO:0010628 | positive regulation of gene expression | 9.52E-08 |
| GO:1903506 | regulation of nucleic acid-templated transcription | 9.14E-08 |
| GO:0006355 | regulation of transcription, DNA-templated | 8.95E-08 |
| GO:2001141 | regulation of RNA biosynthetic process | 6.77E-08 |
| GO:0009058 | biosynthetic process | 6.38E-08 |
| GO:0032879 | regulation of localization | 2.76E-08 |
| GO:0044085 | cellular component biogenesis | 2.18E-08 |
| GO:0022607 | cellular component assembly | 1.35E-08 |
| GO:0042221 | response to chemical | 1.18E-08 |
| GO:0010556 | regulation of macromolecule biosynthetic process | 1.17E-08 |
| GO:0006950 | response to stress | 1.13E-08 |
| GO:0051252 | regulation of RNA metabolic process | 6.40E-09 |
| GO:0046483 | heterocycle metabolic process | 5.80E-09 |
| GO:2000112 | regulation of cellular macromolecule biosynthetic process | 3.94E-09 |
| GO:0051173 | positive regulation of nitrogen compound metabolic process | 3.48E-09 |
| GO:0048583 | regulation of response to stimulus | 2.30E-09 |
| GO:0006725 | cellular aromatic compound metabolic process | 2.22E-09 |
| GO:0007399 | nervous system development | 1.62E-09 |
| GO:0031325 | positive regulation of cellular metabolic process | 1.03E-09 |
| GO:0019219 | regulation of nucleobase-containing compound metabolic process | 1.00E-09 |
| GO:0031326 | regulation of cellular biosynthetic process | 9.98E-10 |
| GO:0023052 | signaling | 6.33E-10 |
| GO:0034641 | cellular nitrogen compound metabolic process | 4.19E-10 |
| GO:0009889 | regulation of biosynthetic process | 3.81E-10 |
| GO:1901360 | organic cyclic compound metabolic process | 3.47E-10 |
| GO:0006996 | organelle organization | 1.12E-10 |
| GO:0010604 | positive regulation of macromolecule metabolic process | 9.91E-11 |
| GO:0007154 | cell communication | 6.24E-11 |
| GO:1901564 | organonitrogen compound metabolic process | 5.28E-11 |
| GO:0010468 | regulation of gene expression | 2.64E-11 |
| GO:0065008 | regulation of biological quality | 2.46E-11 |
| GO:0006810 | Transport | 1.75E-11 |
| GO:0051234 | establishment of localization | 9.61E-12 |
| GO:0009893 | positive regulation of metabolic process | 4.75E-12 |
| GO:0044260 | cellular macromolecule metabolic process | 4.70E-12 |
| GO:0009653 | anatomical structure morphogenesis | 4.67E-13 |
| GO:0051716 | cellular response to stimulus | 3.13E-13 |
| GO:0048513 | animal organ development | 1.86E-14 |
| GO:0048523 | negative regulation of cellular process | 1.50E-14 |
| GO:0030154 | cell differentiation | 4.03E-16 |
| GO:0048869 | cellular developmental process | 2.94E-16 |
| GO:0051179 | localization | 2.48E-16 |
| GO:0080090 | regulation of primary metabolic process | 1.60E-16 |
| GO:0051171 | regulation of nitrogen compound metabolic process | 1.10E-16 |
| GO:0048519 | negative regulation of biological process | 4.22E-18 |
| GO:0060255 | regulation of macromolecule metabolic process | 2.62E-18 |
| GO:0031323 | regulation of cellular metabolic process | 2.40E-18 |
| GO:0043170 | macromolecule metabolic process | 1.69E-18 |
| GO:0048522 | positive regulation of cellular process | 3.22E-21 |
| GO:0048731 | system development | 1.18E-21 |
| GO:0019222 | regulation of metabolic process | 4.48E-22 |
| GO:0071840 | cellular component organization or biogenesis | 4.40E-23 |
| GO:0016043 | cellular component organization | 3.15E-23 |
| GO:0006807 | nitrogen compound metabolic process | 2.83E-24 |
| GO:0050896 | response to stimulus | 6.13E-25 |
| GO:0048518 | positive regulation of biological process | 2.98E-25 |
| GO:0044237 | cellular metabolic process | 4.27E-27 |
| GO:0007275 | multicellular organism development | 2.51E-27 |
| GO:0044238 | primary metabolic process | 7.38E-29 |
| GO:0048856 | anatomical structure development | 6.20E-31 |
| GO:0071704 | organic substance metabolic process | 1.52E-32 |
| GO:0032502 | developmental process | 1.49E-32 |
| GO:0008152 | metabolic process | 6.05E-34 |
| GO:0032501 | multicellular organismal process | 1.33E-34 |
| GO:0050794 | regulation of cellular process | 1.35E-44 |
| GO:0050789 | regulation of biological process | 2.26E-55 |
| GO:0065007 | biological regulation | 3.02E-62 |
| GO:0009987 | cellular process | 1.67E-99 |
